# Whole genome resequencing data enables a targeted SNP panel for conservation and aquaculture of *Oreochromis* cichlid fishes

**DOI:** 10.1101/2021.03.24.436760

**Authors:** A Ciezarek, AGP Ford, GJ Etherington, N Kasozi, M Malinsky, T Mehta, L Penso-Dolfin, BP Ngatunga, A Shechonge, R Tamatamah, W Haerty, F Di Palma, MJ Genner, GF Turner

**Author notes:** Authors contributed equally (A Ciezarek, AGP Ford).

## Abstract

Cichlid fish of the genus *Oreochromis* form the basis of the global tilapia aquaculture and fisheries industry. Non-native farmed tilapia populations are known to be widely distributed across Africa and to hybridize with native *Oreochromis* species. However, many species are difficult to distinguish morphologically, hampering attempts to maintain good quality farmed strains or to identify pure populations of native species. Here, we describe the development of a single nucleotide polymorphism (SNP) genotyping panel from whole-genome resequencing data that enables targeted species identification in Tanzania. We demonstrate that an optimized panel of 96 genome-wide SNPs based on F_ST_ outliers performs comparably to whole genome resequencing in distinguishing species and identifying hybrids. We also show this panel outperforms microsatellite-based and phenotype-based classification methods. Case studies indicate several locations where introduced aquaculture species have become established in the wild, threatening native *Oreochromis* species. The novel SNP markers identified here represent an important resource for assessing broodstock purity and helping to conserve unique endemic biodiversity, and in addition potentially for assessing broodstock purity in hatcheries.

## Introduction

Global aquaculture production has increased rapidly in recent decades. Continued expansion is particularly important in Africa, where rapid human population growth over this century will stress food production systems (FAO, 2020). Tilapia, cichlid fish of the genus *Oreochromis*, native to Africa and the Middle East, have been a key part of the expansion of tropical aquaculture, accounting for 5.5 million of the global total of 47 million tonnes of inland finfish aquaculture production in 2018 (FAO, 2020). However, farmed populations have frequently colonized water catchments where they are not native, both due to deliberate introductions and accidental escape from fish farms (Shechonge, Ngatunga, Bradbeer, et al., 2019). This has threatened native species through ecological competition, habitat alteration and hybridization (Bbole et al., 2014; Canonico et al., 2005; Deines et al., 2014; Firmat et al., 2013; Macaranas et al., 1986; Ndiwa et al., 2014; Waiswa Mwanja et al., 2012). At present native *Oreochromis* species are poorly characterized, and their conservation could benefit from the identification of purebred populations for protection. Such safeguarding of the wild relatives of farmed species would also protect unique genetic resources that could be used to enhance traits in cultured *Oreochromis* strains (Macaranas et al., 1986; Thodesen et al., 2013).

Tanzania, a hotspot of natural diversity for tilapia species, has eight fully endemic *Oreochromis* species (*O. amphimelas, O. chungruruensis, O. karomo, O. korogwe, O. latilabris, O. ndalalani, O. rukwaensis, O. urolepis*). It also has an additional 12 species that are endemic to catchments shared with neighboring countries (*O. alcalicus, O. esculentus, O. girigan, O. hunteri, O. jipe, O. karongae, O. lidole, O. malagarasi, O. pangani, O. squamipinnis, O. tanganicae, O. variabilis*). Several of these species are adapted to unique environmental conditions, such as elevated temperatures, salinity, and pH (Ford et al., 2019; Trewavas, 1983). In addition, although Tanzania hosts a native population of *O. niloticus* indigenous to Lake Tanganyika (Shechonge, Ngatunga, Tamatamah, et al., 2019), non-native farmed populations, largely sourced from Lake Victoria, have been widely distributed across the country (Kajungiro et al., 2019; Moses et al., 2020). The spread of *O. niloticus* has been accompanied by *O. leucostictus*, another species present in Lake Victoria (Bradbeer et al., 2019; Shechonge, Ngatunga, Bradbeer, et al., 2019; Shechonge et al., 2018). The Lake Victoria populations of both *O. niloticus* and *O. leucostictus* were themselves introduced from the Nile system, mostly likely Lake Albert, during the 1950s (Balirwa, 1988).

Nile tilapia (*O. niloticus*), in particular, is becoming established across Africa outside of its natural range, including in South Africa (D’Amato et al., 2007), Zambia (Deines et al., 2014), Zimbabwe (Marufu & Chifamba, 2013), the Democratic Republic of Congo (Goudswaard et al., 2002; Mamonekene & Stiassny, 2012), Kenya (Angienda et al., 2011), as well as Tanzania (Shechonge, Ngatunga, Bradbeer, et al., 2019), with reports of either replacement of the native species or extensive introgressive hybridization (Bradbeer et al., 2019; Shechonge, Ngatunga, Bradbeer, et al., 2019; Shechonge et al., 2018). There is also evidence of parasite transmission from introduced tilapia species to native species (Jorissen et al., 2020). Despite this, intentional movement and stocking of tilapia species into natural water bodies continues in many regions of Africa (Genner et al., 2013).

Several studies have shown that diagnosis of *Oreochromis* hybrids purely based on phenotypic traits of colour or morphology is unreliable (Bbole et al., 2014). Genetic analysis is therefore necessary to determine if introduced and native strains are interbreeding, as well as assessing broodstock purity in commercial aquaculture centres. Mitochondrial DNA has proved insufficient for species diagnosis and resolution of many tilapia species due to recent hybridization (Mojekwu et al. 2021). On the other hand, recent studies have shown the utility of nuclear single nucleotide polymorphism (SNP) data for species and strain diagnosis between species of *Oreochromis* (Syaifudin et al., 2019) and between strains of *O. niloticus* (Lind et al., 2019). Meanwhile, high-throughput sequencing has proved useful in the development of population-specific or species-diagnostic SNP panels for several commercially important fisheries species, including Atlantic salmon (Campbell & Narum, 2011; Larson et al., 2014), European herring (Helyar et al., 2012), Pacific lamprey (Hess et al., 2015) and white bass (Zhao et al., 2019).

Here, we use whole genome resequencing aligned to the Nile tilapia genome assembly (Brawand et al., 2014; Conte et al., 2019) to identify species-informative SNPs distinguishing native and introduced tilapia species important to aquaculture in Tanzania. We also use the SNP panel to identify hybrids in wild populations showing improvement on previous phenotypic and molecular methods for species identification.

## Materials and Methods

### Sample collection

Samples of 12 *Oreochromis* species in Tanzania were collected by experimental seine netting or purchasing fish directly from fish markets or landing sites. Voucher specimens were stored in 80% ethanol (with photographs taken to record live coloration before preservation), and fin clips for genetic analysis were preserved in 96-100% ethanol or DMSO salt buffer.

Specimen ID and sample collection localities are detailed in Table S1. Specimens were identified to species level in the field based on phenotype following diagnostic criteria (Genner et al., 2018). Putative purebred or hybrid status was estimated by a consensus of experienced field researchers from colour photos taken in the field upon collection. Phenotypically identified hybrids either had intermediate phenotypic traits, or discordant combinations of traits typical of purebreds. Species assignments from the genetic data were also compared against this phenotypic ID.

### SNP panel design material

A set of 25 reference individuals from four species were used to identify optimal SNPs for the panel. These reference individuals were from putatively pure populations of *O. urolepis* (n=10) from the Lower Wami and Rufiji rivers, *O. niloticus* (n=6) and *O. leucostictus* (n=6) from Lake Albert and Lake Victoria, as well as samples from *O. shiranus* (n=3; two from Lituhu and one from aquarium stock at Bangor University) (Figure 1; Table S1). These individuals were classified as reference based on a lack of hybrids or other species in the sampling location and confident morphological identification. A further 75 individuals from an additional eight species were included for joint genotyping, as well as testing the ability of the SNP panel to distinguish species not involved in panel design. Collectively these 100 individuals are referred to herein as the “panel design dataset”

**Figure 1.**
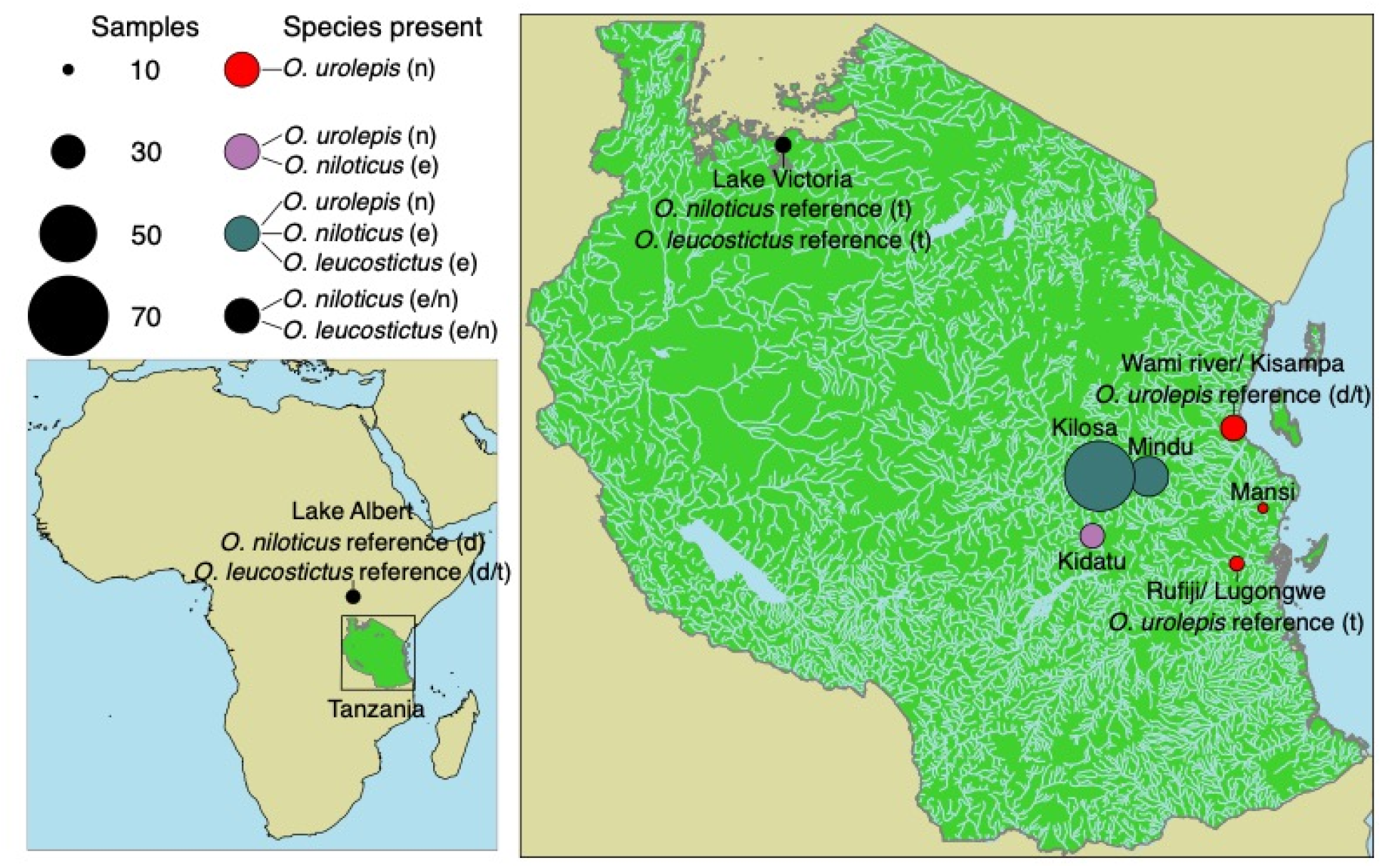
Sample locations for the three focal species within Tanzania (right panel) and Lake Albert (Uganda; bottom left). Abbreviations in ‘Species present’: n: native; e: exotic. Abbreviations in reference notations on map: d: genome-wide sequencing used for design of SNP array; t: test individuals sequenced using SNP array, used as reference for assigning species. Shapefiles sourced from the ArcGIS Hub (continental boundaries), the ICPAC GeoPortal (Tanzania rivers) and the Humanitarian Data Exchange (Tanzania boundary).

### SNP panel performance test material

To study the performance of the SNP panel in hybrid identification, we analyzed samples collected during Feb 2015 and May 2016 from i) the Mindu Dam on the Ruvu River near Morogoro (ii) sites near Kilosa on the Wami catchment, iii) the Kidatu reservoir on the Great Ruaha River – a tributary of the Rufiji system, and iv) sites near Utete on the floodplain of the lower Rufiji River, including the oxbow Lake Lugongwe (Figure 1; Table S1). This is subsequently referred to herein as the “panel test dataset”. The native species at all four locations is *O. urolepis*, also known in aquaculture literature as *O. urolepis hornorum* or *O. hornorum* (Trewavas 1983). Previous work indicates that the Lugongwe site contains pure *O. urolepis*, while the introduced *O. niloticus* is established at Kidatu, and both *O. niloticus* and the non-native *O. leucostictus* are present at Mindu and Kilosa (Shechonge et al., 2018). Microsatellite analysis suggested that hybridization was occurring between both introduced species and *O. urolepis* at Mindu and between *O. niloticus* and *O. urolepis* at Kidatu (Shechonge et al., 2018). Reference individuals were included within the panel test dataset (not the same reference individuals as in the panel design dataset). Specifically, six *O. niloticus*, eight *O. leucostictus* and 14 *O. urolepis* individuals were identified based on a lack of hybrids or other species in the sampling location and confident morphological identification.

### DNA extraction, whole genome resequencing, read mapping and variant calling

DNA for whole genome resequencing was extracted from fin clips using a PureLink^®^ Genomic DNA extraction kit (Life Technologies). DNA extractions for SNP genotyping were processed using the PureLink Genomic DNA kit or a high-salt extraction protocol. Genomic libraries for paired-end sequencing on the Illumina HiSeq 2500 machine were prepared according to Illumina TruSeq HT protocol to obtain paired-end reads, by the Earlham Institute Genomic Pipelines team. For the 100 individuals in the panel design dataset, low coverage sequencing (target 5X mean depth per sample) were sequenced on the Illumina HiSeq 2500 using version 4 chemistry and a 125bp paired-end reads. For the 35 samples in the panel test dataset, sequencing was instead on the Illumina NovaSeq 6000, using 150bp paired-end reads, with an average mean depth of 9x per sample. Raw reads will be made available in the European Nucleotide Archive upon acceptance for publication.

For the panel design dataset, quality analysis of raw reads was carried out using fastQC (v0.11.1) (Andrews, 2010). Alignment and duplicate removal were conducted using a local (Earlham Institute) instance of the Galaxy platform (Blankenberg et al., 2010; Giardine et al., 2005; Goecks et al., 2010). Low coverage reads were all aligned to the *Oreochromis niloticus* reference genome [consisting of the NCBI Orenil1.1 genome version GCA_000188235.2 (Brawand et al., 2014), concatenated with the NCBI mitochondrial genome GU238433.1], using the default settings of BWA-MEM (Galaxy tool version 0.7.12.1) (Li, 2013). Duplicates were removed using the samtools (Galaxy tool version 2.0) rmdup tool (Li et al., 2009). Local realignment around indels was performed per sample using the IndelRealigner tool from software package GATK v 3.5.0 (McKenna et al., 2010). A reference sequence dictionary was created for the reference file using PicardTools v.1.140 (http://broadinstitute.github.io/picard), and the index files for the reference and aligned bam files were created using samtools (v.1.3) faidx and samtools index.

SNP and short indel variants were called against the reference genome using GATK v 3.5.0 Haplotypecaller, using the options -ERC GVCF to output gvcf format, and –minPruning and –minDanglingBranchLength parameters set to 1, to account for the low levels of coverage in the resequencing dataset. Variants were called using a sequence dataset for 100 individuals including pure *Oreochromis* species and putative hybrids. Variant evaluation was performed using PicardTools Collect Variant Calling Metrics function. Output variants were separated by SNP/indel and nuclear/mitochondrial scaffolds, and thereafter analysed separately. Indel files were used to mask indels and sites within 5 bp of indels in the gvcf files,but were otherwise not included in analysis. Variant filtration was performed using GATKs VariantFiltration tool using the following hard filters: QD (QualbyDepth): < 2.0; FS (FisherStrandBias) > 20.0; SOR (StrandOddsRatio) > 4.0; MQ (RMSMappingQuality) > 40.0; MQRS (MappingQualityRankSumTest) < −2.5; RPRS (ReadPosRankSumTest) < −2.0.

For the panel test whole-genome resequence data, variants were called against a newer version of the *Oreochromis niloticus* reference genome (Conte et al., 2019; GCF_001858045.2), not including any of the individuals used to design the panel (Table S2). SNPs were called as for the SNP array design whole-genome calls, except for slightly different filtering parameters (sites excluded with Quality-by-depth < 2, FS > 60, MQ < 40.0, MQRankSum < −12.5, ReadPosRankSum < −8 or total depth less than 100 or greater than 3000). Unlike the SNP array design genotype calls, the galaxy toolkit was not used for this dataset, and different versions of bwa (v0.7.17) and samtools (v1.10) were used. Using bcftools (v1.10.2), biallelic SNPs with a minor-allele count of at least three were extracted and pruned for missing taxa less than 50% and linkage using the prune function, removing SNPs with R^2^ greater than 0.6 over 50kb windows.

### Identification of SNPs for the panel

Biallelic nuclear SNPs from the panel design SNP set (only aligned to linkage groups and excluding those mapped to unplaced scaffolds) were extracted, and the dataset was filtered to include only the 25 reference individuals. Vcftools (v0.1.13) (Danecek et al., 2011) was used to calculate pairwise F_ST_ values (-vcf-weir-pop) between each of the reference species groups (*O. urolepis* n=10; *O. niloticus* n=6; *O. leucostictus* n=6; *O. shiranus* n=3). The SNP set was filtered to include pairwise F_ST_ values >0.9 for at least three of the six pairwise reference population comparisons. The SNP list was further filtered by imposing a minimum distance of 2mn bp between SNPs and ensuring an even spread of high FM_ST_ comparisons across all linkage groups (Table S3).

To examine how the SNP set performs in resolving species, the SNPs included in the panel were extracted from the vcf file for all 100 individuals from all 12 study species. Principal Components Analysis (PCA) was then carried out using SNPRelate (Zheng et al., 2012) in R (R Core Team 2019) and plotted using ggplot2 (Wickham, 2016). A neighbor-joining tree was also inferred using the ‘ape’ package in R (Paradis et al., 2004), using genetic distances calculated using VCF2Dis (https://github.com/BGI-shenzhen/VCF2Dis; accessed December 2020).

As the initial analysis and SNP panel design was conducted using an older version of the *O. niloticus* genome assembly, coordinates were subsequently converted to the latest version of the *O. niloticus* reference genome (Conte et al., 2019; GCF_001858045.2) using the NCBI remap tool (https://www.ncbi.nlm.nih.gov/genome/tools/remap: accessed August 2020). Coordinates for both versions of the reference genome are given in Table S4.

### SNP panel sequencing

The selected SNPs were prepared for panel design by extracting a 50bp flanking sequence either side of the SNP locus from the reference genome assembly. Agena Bioscience^®^ (San Diego, California) SNP genotyping of the selected 120 SNPs was performed at the Wellcome Sanger Institute for n=164 samples (see Table S1). This included the remaining reference individuals not used for SNP panel design as well as all the test individuals. Primer and probe sequences for the genotyping are given in Table S4.

### SNP panel downsampling

To test how many SNPs from the panel are necessary to accurately detect hybrids and assign species, we generated 100 replicates of random subsets of 10, 20, 30, 40, 50, 60, 70, 80, 90, 96, 100 and 110 SNPs. We also tested an optimum set of 96 SNPs, according to principal component loading scores, which we calculated for each SNP using PLINK (v1.90) (Purcell et al., 2007). We selected the 96 SNPs with the highest absolute loading values for PC1 and PC2 combined. We tested the log-likelihood of fastSTRUCTURE (v1.0.0) runs from *K*=1-12 for each replicate, the number of hybrids (individuals with no ancestry component > 80% identified by fastSTRUCTURE, following Shechonge et al., (2018), for each replicate, and the variability between replicates with the same number of SNPs in ancestry component of each identified hybrid. The optimal 96 SNP set was also compared with the full SNP set using fastSTRUCTURE from *K*=1-12.

### SNP panel comparison to other datasets

As genotyping is frequently performed in 96-well plates, the optimum panel of 96 SNPs (described above) was compared to the full 120 SNPs to compare performance. Results of the 96 SNP panel genotyping were also compared to whole genome resequencing analysis, existing published microsatellite data (Shechonge et al., 2018) and the morphological identification. Microsatellite data was available for 54 of the same individuals used for the SNP panel, and whole genome data was available for 35 of the same individuals used for the SNP panel (see Table S1). No individuals had data available for all three comparisons. The full genome test individual dataset was as described earlier.

### Population structure and hybrid detection

For the 96 SNP panel and full genome test individual datasets, we performed a Bayesian clustering analysis in the program fastSTRUCTURE (v1.0), running the main algorithm with *K=1-* 12. The optimal *K* value was chosen using the ChooseK.py script within fastSTRUCTURE.

For the microsatellite dataset, STRUCTURE (v2.3.4) (Pritchard et al., 2000) was run with 500,000 iterations, following 250,000 burn-in iterations. Following (Shechonge et al., 2018), prior cluster assignments (using *LOCPRIOR*) were used, identified using the find.clusters algorithm within the R package adegenet (Jombart, 2008), retaining 20 PCA axes and using 1000 iterations. Three clusters (*K*=3) were utilized, corresponding to the three sampled species. Ten independent runs carried out at *K*=3, with the run with the lowest log likelihood utilized to compare to other datasets. The find.clusters algorithm of adegenet was separately run without specifying the number of clusters, to check if the optimal number of clusters according to BIC score differed from the number of species. Additionally, the analysis of microsatellite data was repeated without prior assignments, with 10 replicates for each value of *K*=1-7. The web version of STRUCTURE HARVESTER (Earl & vonHoldt, 2012) was used to infer the most likely value of *K* using Δ*K* (Evanno et al., 2005).

Hierarchical clustering results are influenced by the numbers of individuals sampled in each population (Puechmaille, 2016). To prevent this being a confounding factor for the 96 SNP set versus microsatellite and 96 SNP set versus genome-wide comparisons, the 96 SNP set was pruned to only include the relevant individuals for both comparisons. On each of these subsampled datasets, three independent runs of fastSTRUCTURE were carried out for each value of *K* between 1 and 12. Therefore three separate fastSTRUCTURE analyses were performed for the 96 SNP set; one comprising all individuals, one with only the same individuals as in the microsatellite dataset and another with only the same individuals as the genome-wide dataset.

For each of these analyses, results from one run with the optimal *K* value were used to assign individuals to species. For the microsatellite analysis with *LOCPRIOR* assignments, the run of *K*=3 with the best log-likelihood score was used. The reference individuals were used to classify ancestry components to species. For the test specimens, a threshold of 80% ancestry component was used to designate individuals to a species (Shechonge et al., 2018), and were considered to have a significant ancestry component corresponding to a species if they had at least 20% ancestry component corresponding to it. For example, if an individual had an ancestry component of 67% corresponding the ancestry component found in the reference *O. leucostictus* individuals, and an ancestry component of 33% corresponding to the reference *O. urolepis* individuals, it was designated as a *O. leucostictus* x *O. urolepis* hybrid. If it had an ancestry component of 81% corresponding to the *O. leucostictus* reference individuals, and 19% to the *O. urolepis* individuals, it was classified as a *O. leucostictus*.

To further assess the hybrid status of individuals described as hybrid from the fastSTRUCTURE analysis (two ancestry components > 0.2), we used NewHybrids (v1.1) (Anderson & Thompson, 2002). NewHybrids assesses the posterior probability that an individual comes from one of six classes: nonhybrid of either of two populations, F1 hybrid, F2 hybrid or backcross of either of two populations. As NewHybrids assumes that there are only two parental taxa, we analysed separate datasets consisting only of individuals assigned by fastSTRUCTURE to belong to one of two species, or to be a hybrid between the two species. For each two-species comparison, five independent runs were carried out, each with a burn-in length of 50,000 followed by an MCMC length of 100,000. No prior information was used to designate individuals to either population. We then checked whether all runs converged (less than 0.05 difference between maximum and minimum estimates in posterior probability for each category for each individual between the five runs) and took the mean posterior probability for each.

## Results

### SNP panel design

For each of the 100 specimens used to generate the SNP panel, ≥98% of reads aligned to the reference genome, with ≥80% of reads properly paired (Table S2). Following the GATK pipeline and filtering, 29,657,078 biallelic SNPs were called. This was pruned down to 18,590,392 SNPs with only the 25 reference specimens, including only sites located within the 22 linkage groups with missing data from at most one individual. A set of 4,789 SNPs with a pairwise Fst of least 0.9 in three of the six pairwise reference population comparisons was extracted from this set. The 120 SNP set was extracted from these 4,789 SNPs after pruning by distance. These 120 SNPs had an average pairwise Fst of 0.47 across the six pairwise comparisons. All except three of the SNPs had an Fst of 1.0 in at least one pairwise comparison (Table S3). These SNPs were distributed across 22 linkage groups and separated by at least 1 Mbp (Table S4). Two of these SNPs were subsequently discarded as they failed Agena Biosciences QC during assay genotyping, resulting in an array of 118 SNPs.

PCA of the 118 SNP set extracted from the vcf file with all 100 individuals suggested that species could be distinguished, with *O. niloticus, O. leucostictus*, and *O. urolepis* particularly distinct. Purebred representatives of species used in the design of the panel were distinct, but clustered relatively tightly in the space within the first two PC axes. They also largely formed monophyletic clusters in a neighbor-joining tree inferred from the 118 SNPs, with the exclusion of potential hybrid individuals, for example the sample T3D7, which was morphologically identified as *O. urolepis* but clustered with morphologically identified hybrids (Figure 2b).

**Figure 2.**
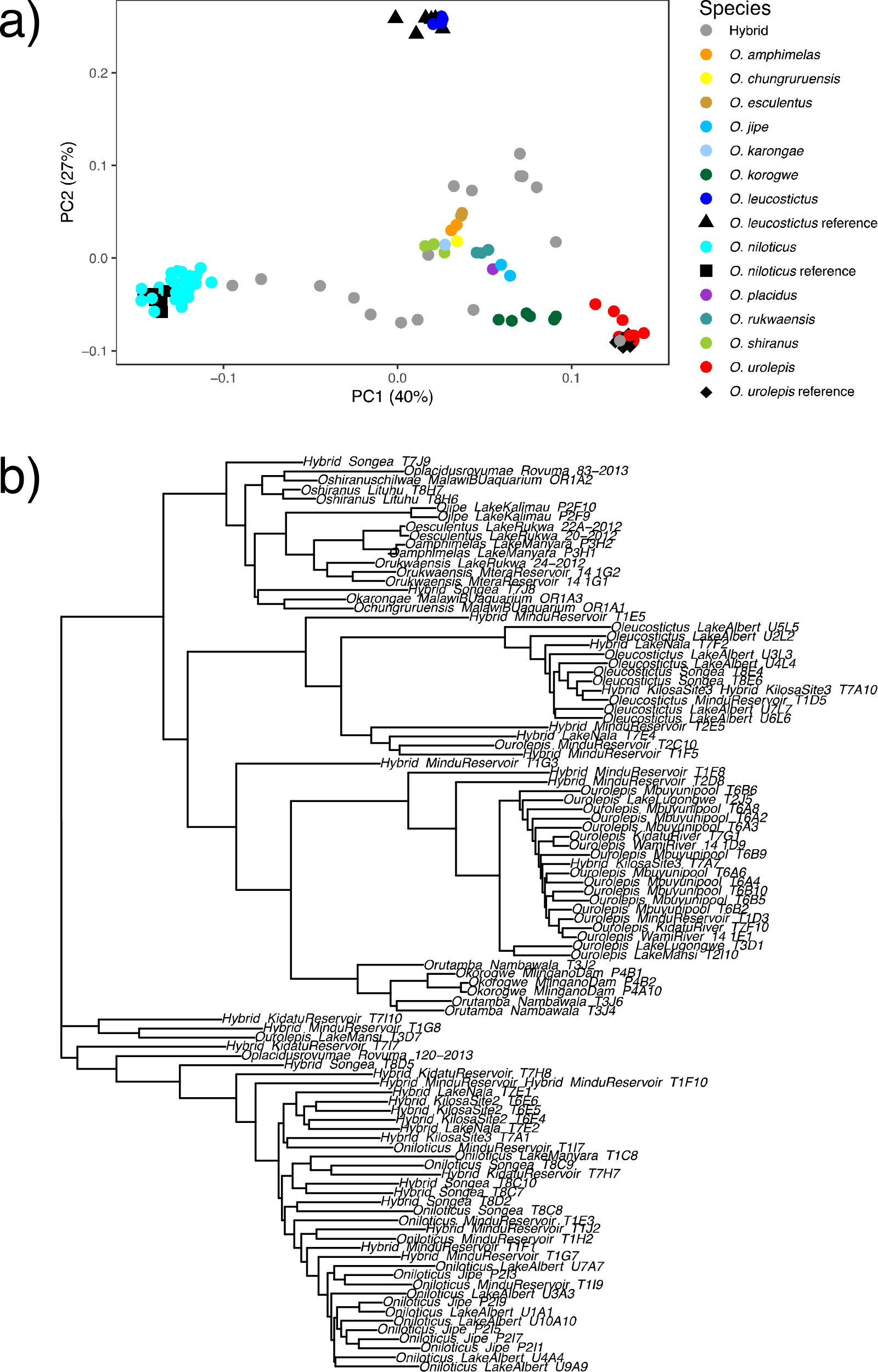
a) PCA of the 118 SNPs, extracted from the full-genome SNP calls from the 100 individuals which were used for initial SNP calling to design the SNP panel. b) neighbor-joining tree of the 118 SNPs from the same 100 individuals. Samples are labelled with their morphological ID, sampling location and sample ID, separated by double underscores.

### The 96 SNP panel is consistent with full-genome data

The 96 SNP set (see Figure S1 for runs from *K*=2-5) and 118 SNP set gave identical classifications for all 164 individuals tested at *K*=3 (the optimal *K* value for both according to chooseK.py) using fastSTRUCTURE (Figure 3a; Table 1).

**Figure 3.**
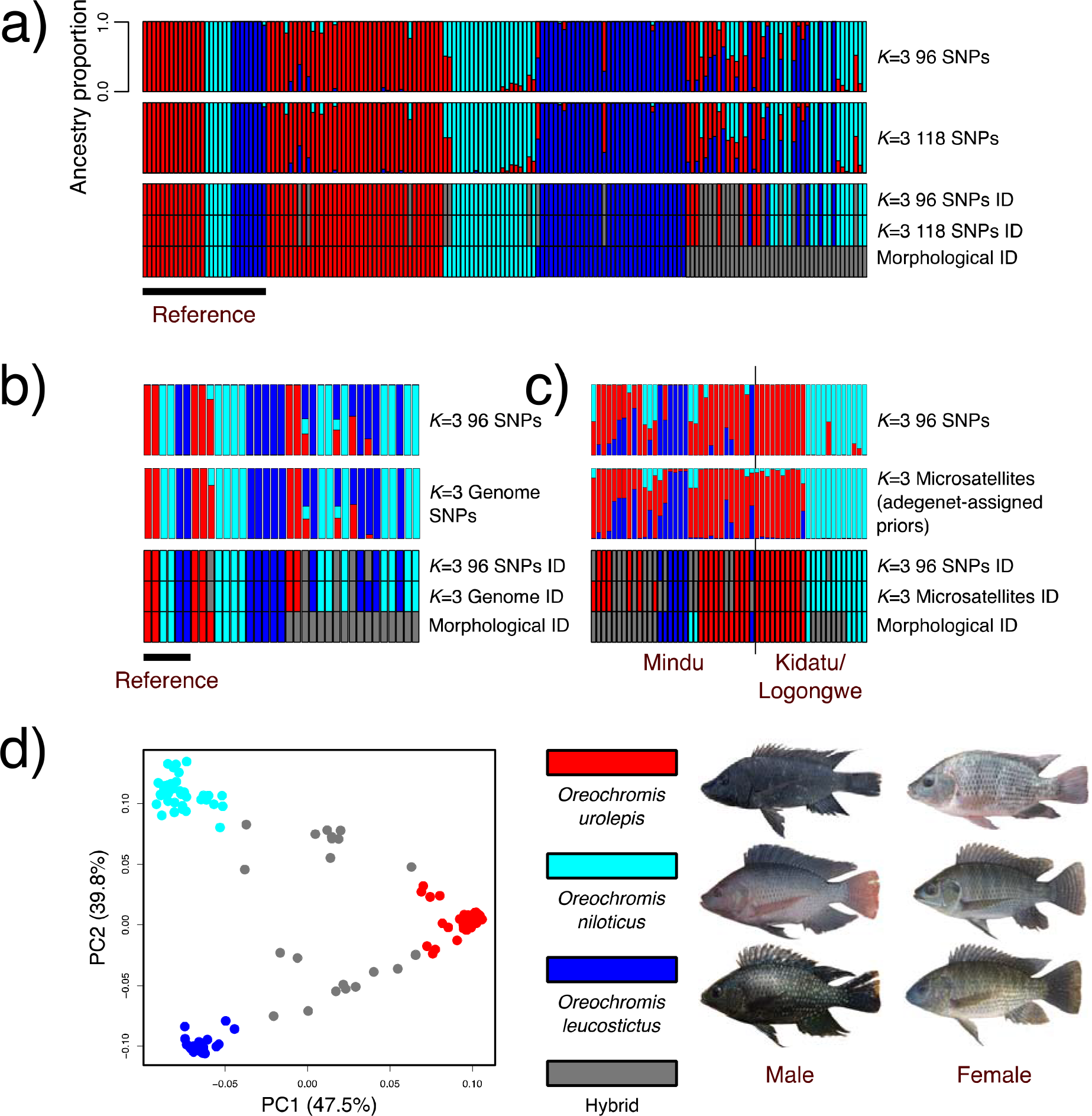
a-c) fastSTRUCTURE analysis, comparison between the 96 SNP set and: a) 118 SNPs; b) genome-wide SNPs; c) microsatellites. d) PCA of the optimal 96 SNP panel, PC1 vs. PC2. The right-hand panel includes representative photographs of mature adults of each species (not to scale).

**Table 1.** Comparison between species assignments between the 96 SNP panel and other datasets.

| Dataset 1 | Dataset 2 | Number of individuals | Number of individuals with the same assignment | Figure |
| --- | --- | --- | --- | --- |
| 96 SNP panel | 118 SNP panel | 164 | 164 (100%) | 3a |
| 96 SNP panel | 18 microsatellite array - prior cluster assignment using adegenet | 54 | 48 (89%) | 3c |
| 96 SNP panel | 1,822,719 genome-wide SNPs | 35 | 34 (97%) | 3b |
| 96 SNP panel | Morphological classification | 164 | 129 (79%) | 3a |
| Morphological classification | 18 microsatellite array - prior cluster assignment using adegenet | 54 | 32 (59%) | 3c |
| Morphological classification | 1,822,719 genome-wide SNPs | 35 | 20 (57%) | 3b |

Following pruning for linkage, 1,822,719 SNPs were used for the genome-wide fastSTRUCTURE analysis. For each of five independent runs *K*=3 was identified as the value to both maximize marginal likelihood and explain the structure in data. Out of the 35 individuals with both genome-wide and 96 SNP data, 34 were identified consistently between the two (Figure 3c; Table 1). The only individual they differed on was identified as *O. leucostictus* in the genome-wide data, whereas the 96 SNP set identified it as a hybrid, with a majority *O. leucostictus* ancestry but also a *O. urolepis* component. Phenotypically, 17 of these individuals were identified as hybrid. However, only 3 were identified as hybrid in the genome-wide data and 4 in the 96 SNP set. Seven of the phenotypic hybrids were identified as *O. niloticus*, two as *O. urolepis* and five as *O. leucostictus* in the 96 SNP set, with four on the genome-wide data.

### The 96 SNP panel outperforms microsatellites

In the STRUCTURE analysis of 18 microsatellites, using the method of Shechonge et al., (2018), where prior assignment of specimens to clusters based on the known number of species (*K*=3), 89% of individuals were given the same assignment as the 96 SNP panel (Table 1; Figure 3c).

However, adegenet incorrectly identified *K*=4 as the optimal number of clusters, according to BIC score. Equally, STRUCTURE analyses with no *a priori* clustering of specimens suggested an optimal *K* value of *K=2*, with another small peak at *K*=5 (Figure S2). In general, assignments without *a priori* information gave an unclear pattern, distinguishing *O. niloticus* and *O. urolepis* from Kidatu and Lugongwe,,but failing to distinguish species in Mindu (Figure S3). The 96 SNP panel, based on the same samples, by contrast, consistently identified *K*=3 as the optimal number of clusters, and reliably distinguished species in Mindu (Figure S3).

### The 96 SNP panel is more accurate than identification based on phenotype

The 96 SNP set assigned all the reference individuals to the same species as the phenotypic ID (Table S5). However, there were some differences in the test individuals of all three species, with three phenotypically identified *O. urolepis*, three phenotypically identified *O. niloticus* and two phenotypically identified *O. leucostictus* being designated as hybrids (all *O. niloticus* x *O. urolepis* or *O. leucostictus* x *O. urolepis*). One phenotypically identified *O. leucostictus* was instead classified as *O. urolepis* by the 96 SNP panel. Many of the phenotypically identified hybrids were instead given pure species classification: six as *O. urolepis*, 14 as *O. niloticus* and six as *O. leucostictus*. Only 14 were identified as hybrid by both phenotype and the 96 SNP panel (Figure 3a).

### Validation of hybrid classification

In total, 96% of fastStructure identifications were corroborated with NewHybrids, with posterior probability > 0.98 (Table S6,7). Two NewHybrids analyses were carried out: one with *O. urolepis* and *O. niloticus* individuals, and individuals identified as hybrid between the two, and one with *O. urolepis* and *O. leucostictus* individuals, and their hybrids. Comparisons were not made between *O. niloticus* and *O. leucostictus*, as no hybrids were identified between the two using fastSTRUCTURE. Six F1 hybrids were identified between *O. urolepis* and *O. niloticus* (posterior probability > 0.95), alongside five *O. urolepis* backcrosses and three *O. niloticus* backcrosses (Table S6). Five F1 hybrids (posterior probability > 0.95) were identified between *O. leucostictus* and *O. urolepis*, alongside one F2 hybrid. Six *O. urolepis* backcrosses were also identified, with three *O. leucostictus* backcrosses (Table S6).

Together, we found evidence of introgression between the native *O. urolepis* and both the invasive *O. leucostictus* and *O. niloticus* in Kilosa, Kidatu and Mindu. We find no evidence of hybrids in Rufiji, Lugongwe or Mansi (Table S1). We found no evidence of any hybrids between *O. niloticus* and *O. leucostictus*. See Table 2 for the numbers of each species and hybrid identified by the 96 SNP panel.

**Table 2.** Number of individuals of each species identified in each sampling location by the 96 SNP panel.

| Location | *O. urolepis*<br>individuals | *O. leucostictus*<br>individuals | *O. niloticus*<br>individuals | Hybrids |
| --- | --- | --- | --- | --- |
| Mindu Dam | 13 | 6 | 0 | 7 *leucostictus* x<br>*urolepis*<br>6 *niloticus* x<br>*urolepis*<br>1 *leucostictus* x<br>*niloticus* x<br>*urolepis* |
| Kilosa | 6 | 32 | 18 | 4 *leucostictus* x<br>*urolepis*<br>1 *niloticus* x<br>*urolepis*<br>1 *leucostictus* x<br>*niloticus* x<br>*urolepis* |
| Kidatu resevoir | 5 | 0 | 14 | 2 *niloticus* x<br>*urolepis* |
| Lake Mansi/<br>Lugongwe/<br>Mindu | 20 | 0 | 0 | 0 |

### Different subsets of at least 80 out of the 118 SNP dataset give consistent results

For the majority of subsample replicates of the 118 SNPs, *K*=3 was identified as the model complexity that maximized marginal likelihood for the majority of these replicates with the following exceptions: *K*=2 was optimal for 6/100 10-SNP sets, *K*=4 was chosen for 1/100 20-SNP sets, 1/100 of the 30-SNP sets and 1/100 of the 70-SNP sets, and *K*=5 was chosen for 1/100 of the 60-SNP sets and 1/100 of the 90-SNP sets. The Model components used to explain structure in the data varied much more between replicates, from 1-11.

Increasing the number of SNPs increased likelihoods, with sharp increases from 10-30 SNPs and more modest increases thereafter (Figure S4a). The number of iterations in which all the reference individuals were correctly classified into populations increased with the number of SNPs, until it reached 100% at 80 SNPs (Figure S4b). It also decreased the number of hybrids identified up to 80 SNPs, after which it then stabilized (Figure S5a). Increasing SNP number also increased the stability of the estimated hybrid ancestry proportion, measured as the variability in the minor ancestry component of a hybrid (Figure S5b). In the 96 SNP iterations, 18 individuals were consistently identified as hybrid in all replicates, whereas 9 were sometimes classified as hybrids. Of these, 5 were identified in fewer than 12 out of 100 replicates, and 4 were identified in at least 78 replicates (Figure S5c,d).

## Discussion

We demonstrate that a reduced panel of 96 genome-wide SNPs performs comparatively well to full genome resequencing in distinguishing species and identifying hybrids of *Oreochromis*. We identify replicate cases where introduced aquaculture species have become established and interbred with native species, including backcrosses as well as F1 and F2 hybrids. We demonstrate that hybridization is persistent in the environment with multi-generation hybrids and backcrossing to parental species.

We found that the ability of a reduced 96-SNP panel to detect hybrids was indistinguishable from the full 118 SNPs that we genotyped. This is likely to be the most cost-efficient panel size, as genotyping is frequently performed in 96-well plates. None of these SNPs overlapped with previously identified species-diagnostic SNPs for *O. niloticus* (Syaifudin et al., 2019), likely because our SNP set was optimized by interspecific rather than intraspecific variation. Our analyses indicate that the reduced 96 SNP panel can accurately identify the hybrids between *Oreochromis* species tested, including introgression from the invasive *O. niloticus* and *O. leucostictus* into the native *O. urolepis* (Shechonge, Ngatunga, Bradbeer, et al., 2019). No hybrids were identified between *O. leucostictus* and *O. niloticus* in Tanzania. These two species co-occur in Lake Albert, Uganda, where they are not known to hybridize (Trewavas 1983). It is possible therefore that behavioral, ecological or genomic incompatibilities prevent the two species from hybridizing in populations where they naturally occur, although *O. leucostictus* has been shown to hybridize in Kenya with other subspecies of *O. niloticus*, with which it does not naturally co-exist (Ndiwa et al., 2014).

This detection of introgression between *O. urolepis* and *O. leucostictus*, and between *O. urolepis* and *O. niloticus* was concordant with previous studies using microsatellite data (Shechonge et al., 2018). Our re-analysis with the same set of microsatellites only gave comparable results if *‘LOCPRIOR’* assignments were used based on an initial clustering, which suggests that power was low in the microsatellite analysis due to a small number of markers or samples (Porras-Hurtado et al., 2013). Additionally, using the *‘LOCPRIOR’* required choosing the value of *K* based on sampling (*K*=3), rather than the optimal number according to BIC score of *K*=4. This suggests that an added benefit of the 96 SNP set is that prior assumptions are not necessary to set the appropriate value of *K* when relatively few individuals are sampled, unlike with the microsatellite data. This may be important in cases where an unknown number of test species are sampled, or there is hidden population structure (Porras-Hurtado et al., 2013). The SNP panel would therefore require less thorough sampling to allow accurate species or hybrid assignment.

Notably, our analyses suggested that morphological identification of hybrids was inconsistent with genetic assignments; many individuals phenotypically assigned as hybrids were genetically classified as pure species. This may reflect high phenotypic diversity within species (Table S5), and possibly overlap in characteristics between species, which could be difficult to catalogue, making species identification more challenging. It may also reflect introgression which has been masked by several generations of backcrossing. This would result in small ancestry components for the introgressed species, and incorrect pure species assignment using hierarchical clustering (e.g. STRUCTURE or fastSTRUCTURE) or NewHybrids. Further studies with large sample sizes, thorough population sampling, genomic data and detailed demographic analyses are necessary to identify if this is the case. This would mean introgression has been occurring for several generations, possibly influencing phenotypic variation within species.

Several analyses suggested that the panel of 96 SNPs provides sufficient power to reliably identify these species and hybrids. Importantly, species assignments using the 96 SNP panel were almost identical to those given by genome-wide data (Figure 3b; Table 1). This suggests that adding more SNPs at extra cost would not greatly improve assignment accuracy, and introgression from the invasive species can be detected reliably without the considerable investment of whole-genome or reduced-representation (e.g. RAD-seq) resequencing. SNP panels of similar sizes have proved accurate at detecting hybrid status and introgression between domestic cats and European wildcats (Oliveira et al., 2015), and between farmed and wild Atlantic salmon (Wringe et al., 2019).

Subsamples of the full 118 SNP set further indicated that individuals could be accurately assigned if > 80 SNPs were used. Although hybrids identification was not fully consistent between iterations even when a larger number of SNPs were used, variability in ancestry components between iterations was low (<0.1). This indicates that the individuals which are classified as hybrids in some but not all iterations (Figure S5d) are those with an ancestry component of close to our arbitrary cut-off to define a hybrid. We recommend that any individuals which are close to the cut-off value chosen are further investigated, for example using NewHybrids. The choice of threshold to define a hybrid may also be adjusted depending on the application. For example, if the panel is being used to eliminate hybrids from breeding stock, then it may be necessary to use a stricter threshold to define hybrids. These analyses indicate that 96 SNPs is above the point of diminishing returns for hybrid identification and accurate reference individual identification, meaning that even if some SNPs fail to amplify in some individuals there should still be sufficient power.

It is important to further consider methodological limitations to accurate species and hybrid assignment using the SNP panel. A signal of introgression indicated by hierarchical clustering can be given in the absence of any introgression of one population that has undergone a recent bottleneck, or in the case of ‘ghost’ introgression from an unsampled population (Lawson et al., 2018). Given that introgression was only inferred in some individuals within each population in our analysis, and the general concordance with NewHybrids analysis, it is likely that the signal we are detecting is in fact introgression, rather than a population-level bottleneck. However, this must be a consideration for users applying the SNP panel on other *Oreochromis* species that we have not tested here. The issue of introgression from unsampled taxa is more likely to be confounding in our dataset, given we have only extensively tested three out of the at least 37 species of *Oreochromis* (Ford et al., 2019). As *O. niloticus* and *O. leucostictus* are the only introduced *Oreochromis* species found in the tested water bodies (Shechonge, Ngatunga, Bradbeer, et al., 2019), it is likely that these results do reflect introgression from one of these species. Reassuringly, PCA and a neighbor-joining tree inferred from the 118 SNPs extracted from the individuals with full-genome resequencing suggested that most of the species are distinguishable, particularly the highly invasive *O. niloticus* and *O. leucostictus* (Figure 2), suggesting that introgression from either of these two species would be identifiable. The discriminatory ability of the SNP panel will need to be tested in cases where other Tanzanian native species co-occur with the focal introduced species, as the current SNP panel was not optimized for other species groups. However, even if native species could not convincingly be distinguished, the SNP panel we present will be able to identify introgression from invasive *O. niloticus* and *O. leucostictus*.

Hierarchical clustering results may also be influenced by uneven sampling of populations (Puechmaille, 2016). In the case that only one or two individuals are sequenced from one population in a large dataset, it is unlikely that they will be assigned a distinct cluster, even in the absence of any introgression. This may mean a lot of diversity within the dataset may be missed. Species assignment tools based on network estimation have the potential to identify these ‘outlier individuals’, which do not belong to any of the reference populations (Kuismin et al., 2020). However, it is not clear how they perform in the presence of hybrid individuals. Future studies using this SNP panel will need to prioritize establishing a reference set of individuals belonging to each target species, with a similar number of individuals of each.

We anticipate that our efficient SNP panel will be of use to the aquaculture and conservation genetics communities in assessing broodstock purity, determining hybrid status of wild populations, and identifying populations most in need of conservation resources.

## Acknowledgements

This work was funded by BBSRC award BB/M026736/1 (GFT, MJG, FdP), and Royal Society Leverhulme Trust Africa Awards AA100023 and AA130107 (MJG, BPN, GFT, RT). WH and FDP acknowledge the support of the Biotechnology and Biological Sciences Research Council (BBSRC), part of UK Research and Innovation; this research was funded by the BBSRC Core Strategic Programme Grants BB/CSP1720/1 and its constituent work packages *-* BBS/E/T/000PR9818. AC, WH, FDP were supported through BBSRC GCRF funding (BB/P028098/1). Samples from Tanzania were collected across 2013-2016 under permit numbers 2014-374-ER-2011-103, 2015-83-NA-2011-103, and 2016-293-NA-2011-103. Reference samples from Uganda (Lake Albert) were collected in 2015 under permit number IMP/GEN/2014/06. We thank María Lorena Romero-Martinez for assistance with sample collection and DNA extraction. We thank Richard Durbin and the DNA Pipelines team at the Wellcome Sanger Institute for assistance with designing the SNP panel. This research was supported in part by the NBI Computing infrastructure for Science (CiS) group through use of the CiS high-performance computing cluster for the analysis of the whole genome resequencing.

## Data Accessibility

Whole genome resequencing data will be deposited before publication

SNP datasets: Dryad

## Author Contributions

GFT, MJG, FDP, and MM conceived the study. MJG, GFT, AS, BPN, and RT designed fieldwork and sampling. AGPF, NK, BPN, AS, GFT, RT and MJG conducted or supervised fieldwork, or collected data. AGPF and TM performed laboratory work. AGC, AGPF, GE, LP-D and WH designed and performed the analysis. AGC and AGPF wrote the first draft of the manuscript. All authors commented on and edited the final manuscript.

## Supplementary Tables

Table S1. Sample information, sequencing details and species assignments.

Table S2. Mapping statistics for the whole genome sequence data.

Table S3. Pairwise Fst values for the 120 SNPs.

Table S4. Primer and probe sequences for all validated panels.

Table S5. Morphological identification and photographs of each sample.

Table S6. Individuals with differing assignments between fastSTRUCTURE and NewHybrids.

Table S7. NewHybrids results.

## Supplementary Figures

**Figure S1.**
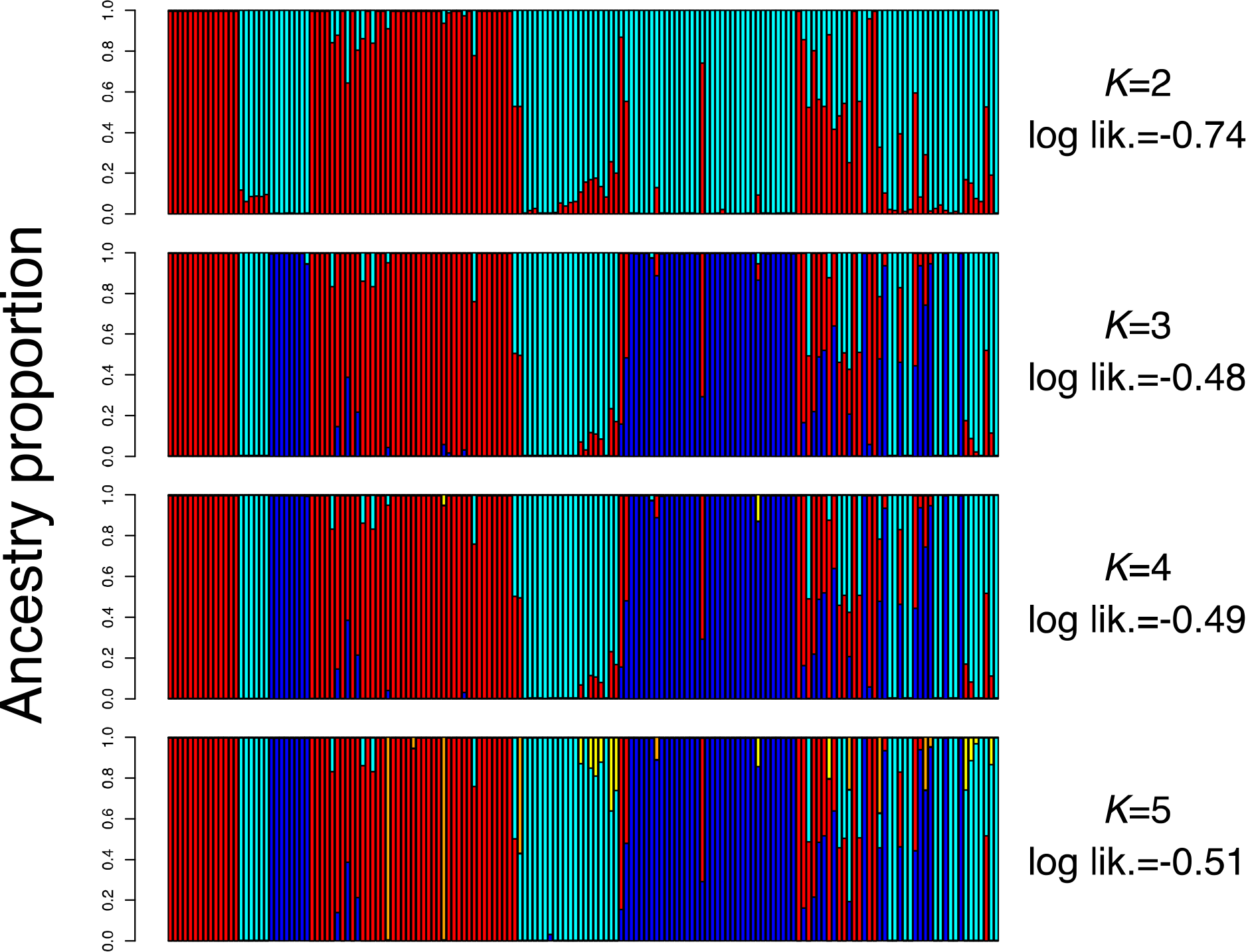
fastSTRUCTURE analysis for all individuals in the 96 SNP panel dataset, from *K*=2 to *K*=5.

**Figure S2.**
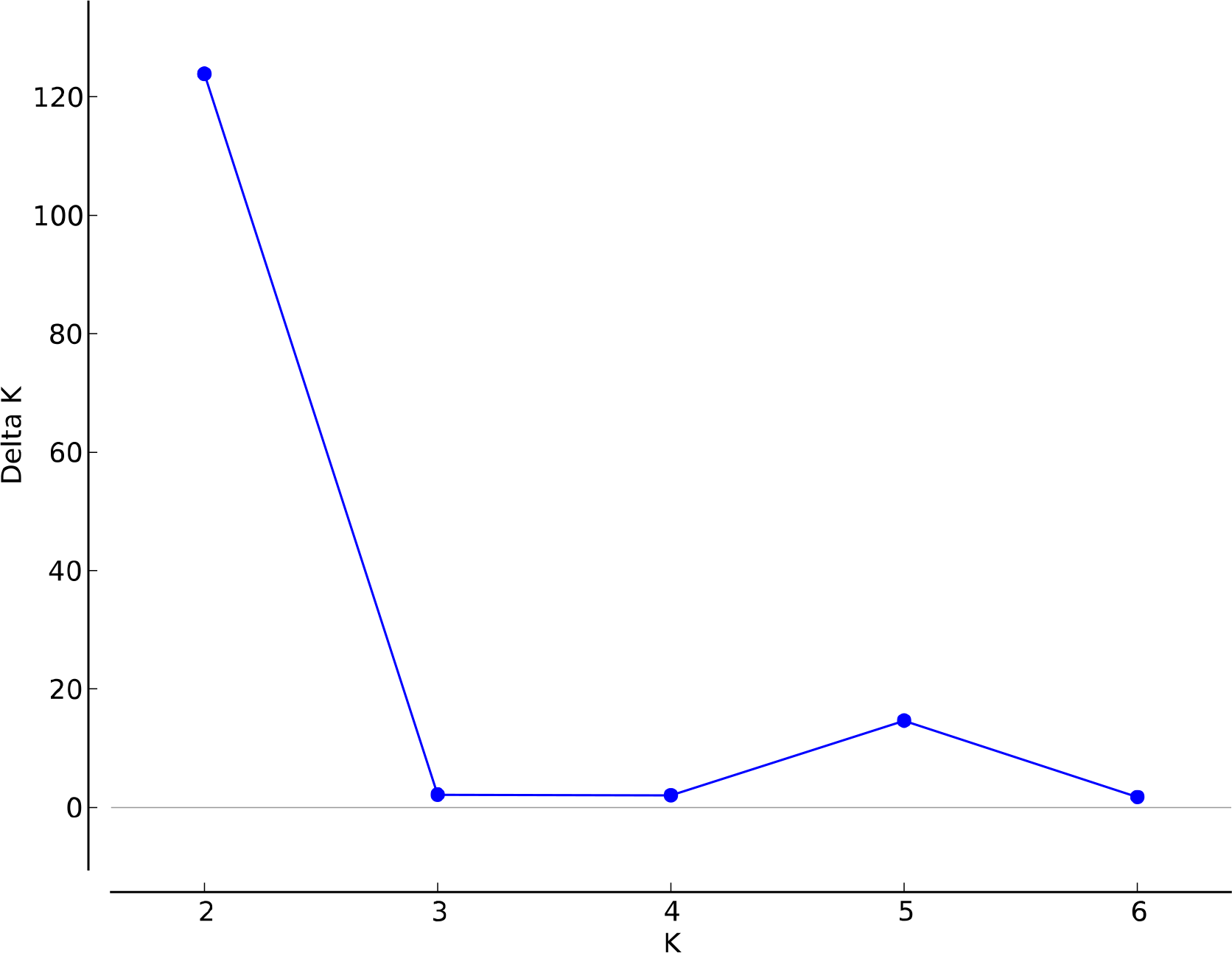
Δ*K* (DeltaK) values for STRUCTURE runs on the microsatellite dataset, without prior assignment, from *K*=2 to *K*=6.

**Figure S3.**
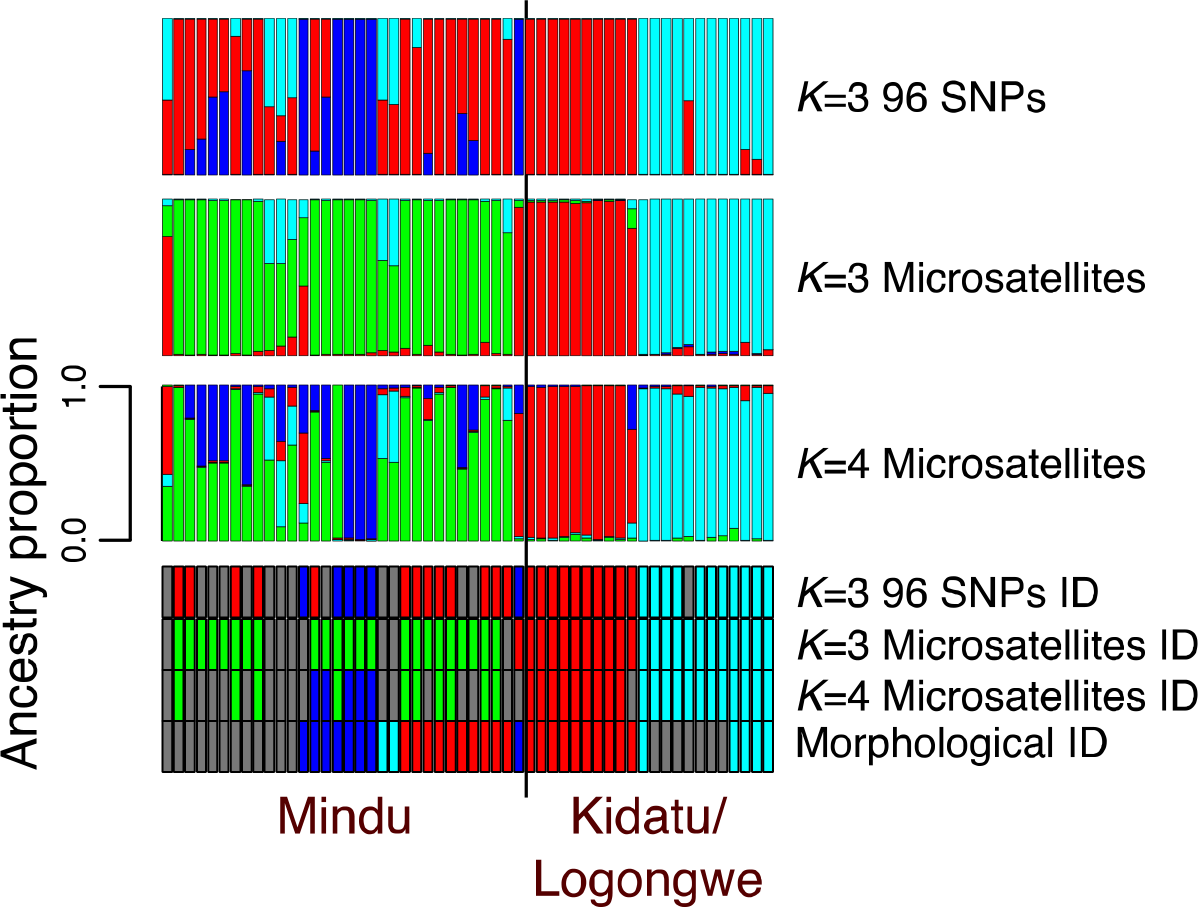
Comparison between species assignment using the 96 SNP panel (top row), and microsatellite analysis without prior assignment at *K*=3 & *K*=4.

**Figure S4.**
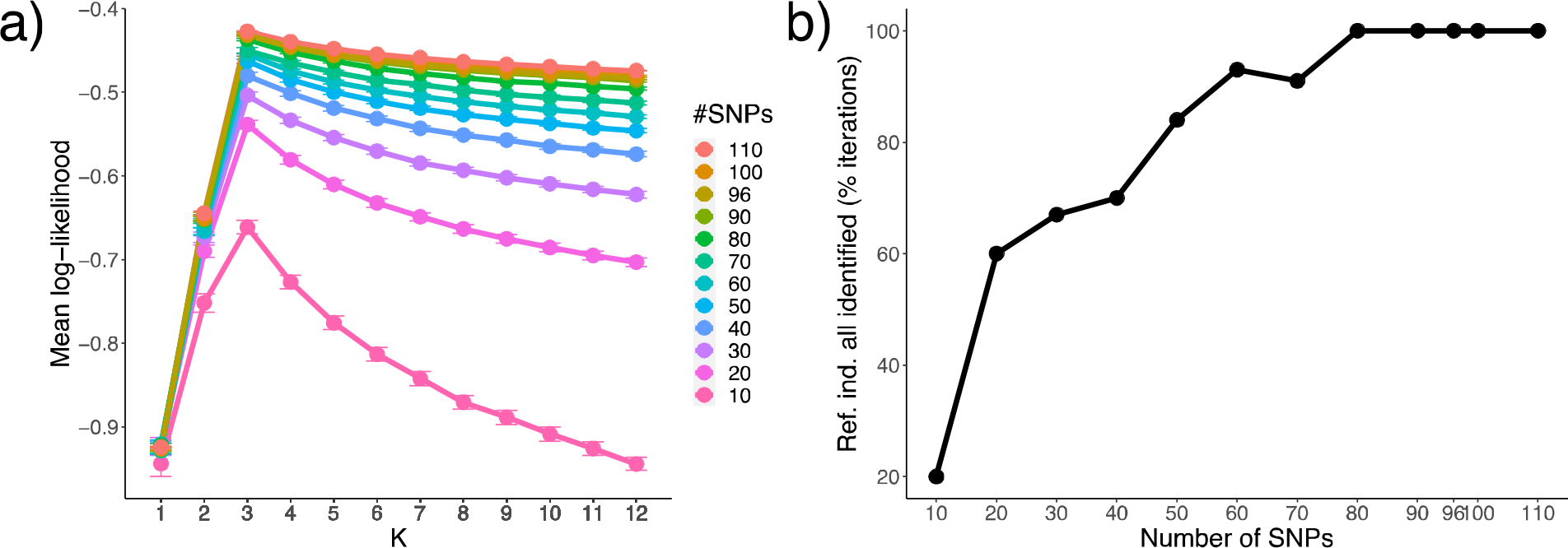
a) Average log-likelihoods for the 100 replicates of each number of sub-sample SNPs. Error bars represent standard error. b) The percentage of replicates for each number of sub-sample where all of the reference individuals were correctly assigned to their species.

**Figure S5.**
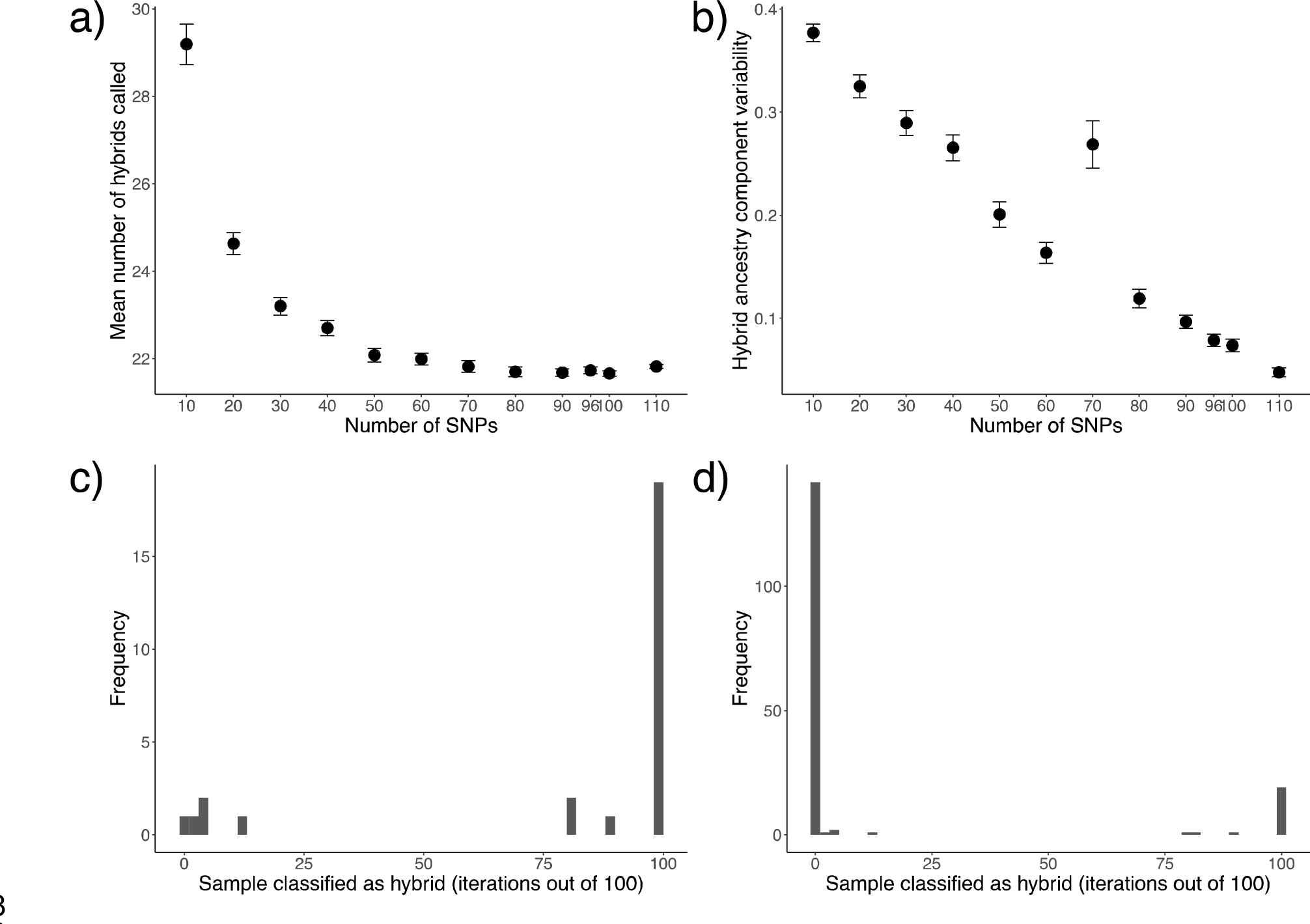
a) The mean number of hybrids called (no ancestry component > 80%) across the 100 replicates of each number of sub-sample SNPs. Error bars represent standard error. b) The mean variability in minor ancestry component between the 100 replicates for each individual identified as hybrid in at least one of these replicates for each number of sub-sample SNPs. Error bars represent standard error. c) Histogram of the frequency at which hybrids were classified as hybrids across the 100 replicates of 96 random SNPs. d) same as c) except that data for individuals never identified as hybrids is added.

